# NAD^+^ precursor treatment prevents cardiomyopathy but disrupts erythroid maturation in mitochondrial progeria

**DOI:** 10.1101/2025.10.02.678456

**Authors:** Nahid A Khan, Kati Ahlqvist, Swagat Pradhan, Juan Landoni, Christopher Jackson, Aleksandra Zhaivoron, Riikka Kivelä, Liliya Euro, Anu Suomalainen

**Affiliations:** Research Programs Unit, Stem Cells and Metabolism, University of Helsinki, 00290 Helsinki, Finland; Institute of Biotechnology, HiLIFE, University of Helsinki, Helsinki, Finland; Department of Biochemistry and Developmental Biology, University of Helsinki, 00290 Helsinki, Finland; Wihuri Research Institute, Helsinki, Finland; Faculty of Sport and Health Sciences, University of Jyväskylä, Jyväskylä, Finland; HUS Diagnostic Center, Helsinki University Hospital,00290 Helsinki, Finland; HiLife, University of Helsinki, 00290 Helsinki, Finland

**Keywords:** NAD^+^ Metabolism, Anemia, Erythropoiesis, Mitochondria, Metabolism, Post mitotic tissues, Progeria

## Abstract

Nicotinamide adenine dinucleotide (NAD^+^) plays a central role in energy metabolism, and its decline is linked to various degenerative diseases. While NAD^+^ restoration holds therapeutic promise, its long term, tissue-specific consequences remain poorly understood. We investigated effects of nicotinamide riboside (NR) supplementation for “mutator” mice manifesting mitochondrial progeria. Our results reveal strikingly divergent outcomes: in proliferative bone marrow, NR-treated mutators show reductive stress with accumulation of NADH/NADPH, altered amino acid, nucleotide, folate levels and impaired heme biosynthesis. In blood, erythrocyte maturation defects are aggravated, exacerbating anemia. Conversely, in postmitotic cardiac tissue, NR enhanced contractility, reduces stress response markers and normalized metabolic profile. These findings indicate that while being beneficial for heart, chronic NAD^+^ boosting can compromise erythrocyte maturation in the context of mitochondrial disease. The data emphasize importance of evaluating systemic effects of NAD^+^ boosting therapies beyond the primary affected tissues and development of tissue-specific metabolic interventions for degenerative diseases.

**Highlights:**

1. Chronic nicotinamide riboside supplementation exacerbates anemia and disrupts erythroid maturation in progeric mice.
2. In proliferative bone marrow cells, NR induces redox imbalance and drives profound metabolic dysregulation.
3. NR suppresses heme biosynthesis and iron transport pathways in the bone marrow.
4. In the heart, NR restores NAD^+^ levels, enhances cardiac function, and reduces metabolic stress.

## Introduction

Nicotinamide adenine dinucleotide (NAD^+^) and its derivatives NADH, NADP and NADPH are central metabolic cofactors essential for maintenance of metabolic homeostasis, with key roles in glycolysis, fatty acid oxidation, and tricarboxylic acid (TCA) cycle^1,2^. Beyond its canonical role in metabolism, NAD^+^ serves as a crucial cofactor and substrate for regulatory proteins such as sirtuins^3^ poly (ADP)-ribose polymerases (PARPs) ^4,5^, and CD38 ^6–8^, thereby regulating mitochondrial metabolism, DNA damage responses, immune function ^9^ and overall cellular resilience. Consequently, NAD metabolite pools profoundly influence cellular function and systemic homeostasis ^10^. NAD-pools are consumed in repair processes, resulting in imbalance between NAD-forms, metabolic remodeling and dysregulated cellular signaling pathways, ultimately compromising physiological fitness and increasing susceptibility to diseases ^1,3,9,11^. Declining NAD^+^ levels have been associated with degenerative ageing related diseases^12^, cancer cachexia^13^, cardiomyopathy^14,15^ and mitochondrial muscle diseases^16^. Supplementation with NAD^+^ precursors (vitamin B3 forms) has shown therapeutic potential and improved mitochondrial function and ameliorated diseases phenotype in mitochondrial myopathy ^17^, improved physical performance ^18–20^, enhancing cardiac function and extending life span in mice ^21–24^.

Similarly, early-phase clinical studies have demonstrated beneficial effects of various NAD^+^ boosters. Treatment with niacin, an NAD^+^ precursor, slowed down progression of mitochondrial myopathy in patients ^16^, nicotinamide riboside (NR) have been shown to modulate NAD metabolome, leading to a reduction in inflammatory cytokines in both serum and cerebrospinal fluid in patients with Parkinson’s disease^25,26^, decreased inflammatory cytokines and reduced ejection fractions in patients with heart failure^27^ and improved muscle mitochondrial biogenesis and satellite cell differentiation in discordant twins^28^. Nicotinamide mononucleotide (NMN) supplementation increased muscle insulin sensitivity in prediabetic women ^29^. Studies in elderly individuals suggested that increasing NAD^+^ levels in muscle decreased the levels of inflammatory cytokines^30^ and improved physical performance^31^. Despite showing therapeutic benefits, some concerning side effects such as mild but consistent decrease in hemoglobin ^16^ and platelet counts^40^ has been observed in patients indicating that knowledge on tissue-specific effects of high-dose NAD precursor treatments are needed.

Given that NAD^+^ metabolism is differentially regulated across cell types and tissues, a more nuanced understanding is needed to optimize therapeutic strategies^9^. Here, we report the impact of long-term NAD^+^ repletion in the progeroid mutator mouse model, which harbors a knock-in mutation in the exonuclease domain of polymerase gamma (POLG) and mitochondrial DNA (mtDNA) mutagenesis, with secondary nuclear DNA damage in proliferating stem and progenitor cells^32^.These mice accumulate random mtDNA mutations in multiple tissues and recapitulates key features of mitochondrial disease such as osteoporosis, hair loss, graying, including cardiomyopathy^33,34^ and life-limiting progressive anemia^35^. We report that NAD^+^ metabolism is compromised in multiple tissues of these mice, and that repletion through NR leads to strikingly divergent tissue-specific responses most notably between the proliferative bone marrow and postmitotic heart. Further, we show that the redox-metabolic consequences of chronic NAD^+^ booster treatment are remarkably different depending on the tissue type. In the proliferative bone marrow cells, it induces significant redox imbalance, characterized by excessive NADH and NADPH accumulation, severe alterations of amino acid levels, glucose, folate, disrupted heme biosynthesis and highly aberrant erythroid maturation, while in the heart NAD^+^ levels increase with sustained cardiac contractility across the mouse lifespan. Our findings highlight the complex and even opposing effects of NAD^+^ booster treatment in solid and proliferative tissues with aging related symptoms.

## Results

### NR treatment exacerbates signs of anemia in aging mice

The phenotypic analysis of mutator mice has been published in detail in a large number of papers, with the first symptom at six months of age being progressive anemia and reduced fertility, followed by progeroid symptoms with osteoporosis, thin skin, gray hair, kyphosis and cardiomyopathy^33-35^. Analysis of NAD^+^ levels in the blood of mutator mice, displayed a significant, age-related decline (Fig. 1A). To assess whether NAD^+^ replenishment could reverse this phenotype, we supplemented 5-month-old mutator mice, still not showing any progeroid features and anemia with nicotinamide riboside (NR; 400 mg/kg/day), a vitamin B3 derivative previously shown to improve muscle metabolism in mitochondrial myopathy model^17^. Hemoglobin levels were measured biweekly for the first five months of treatment and weekly during the sixth month. Although NR supplementation elevated circulating NAD^+^ levels in all treated mice (Fig. 1B), NR-treated mutators exhibited a progressive and unexpected decline in hemoglobin levels relative to untreated mutators and controls (Fig.1C). This was further confirmed by terminal blood analysis, which revealed that mutator mice exhibited lower hemoglobin levels, reduced red blood cell (RBC) counts and RBC percentual amount of blood cells (decreased hematocrit-HCT) compared to control mice. NR treatment exacerbated these hematological abnormalities (Fig. 1D, E, F). Mutator mice also displayed increased variability of RBC size (red cell distribution width; RDW) and distribution of hemoglobin in RBCs (hemoglobin distribution width; HDW). Both signs showed an exacerbating trend after NR treatment (Fig S1 A, B). Still, the mean volume of RBCs (MCV) and their mean erythrocytic hemoglobin concentration (MHCH) remained unchanged (Fig S1D-F). In addition to anemia, mutator mice displayed reduced white blood cell (WBC) counts compared to wild-type (WT) mice which remains unchanged after NR (Fig. 1G). However, the platelet counts showed decrease in both wild-type and mutator mice following NR treatment (Fig S1C). These findings indicated that the effect of NR was especially severe for RBCs and platelets, while white blood cells were not affected.

**Figure 1.**
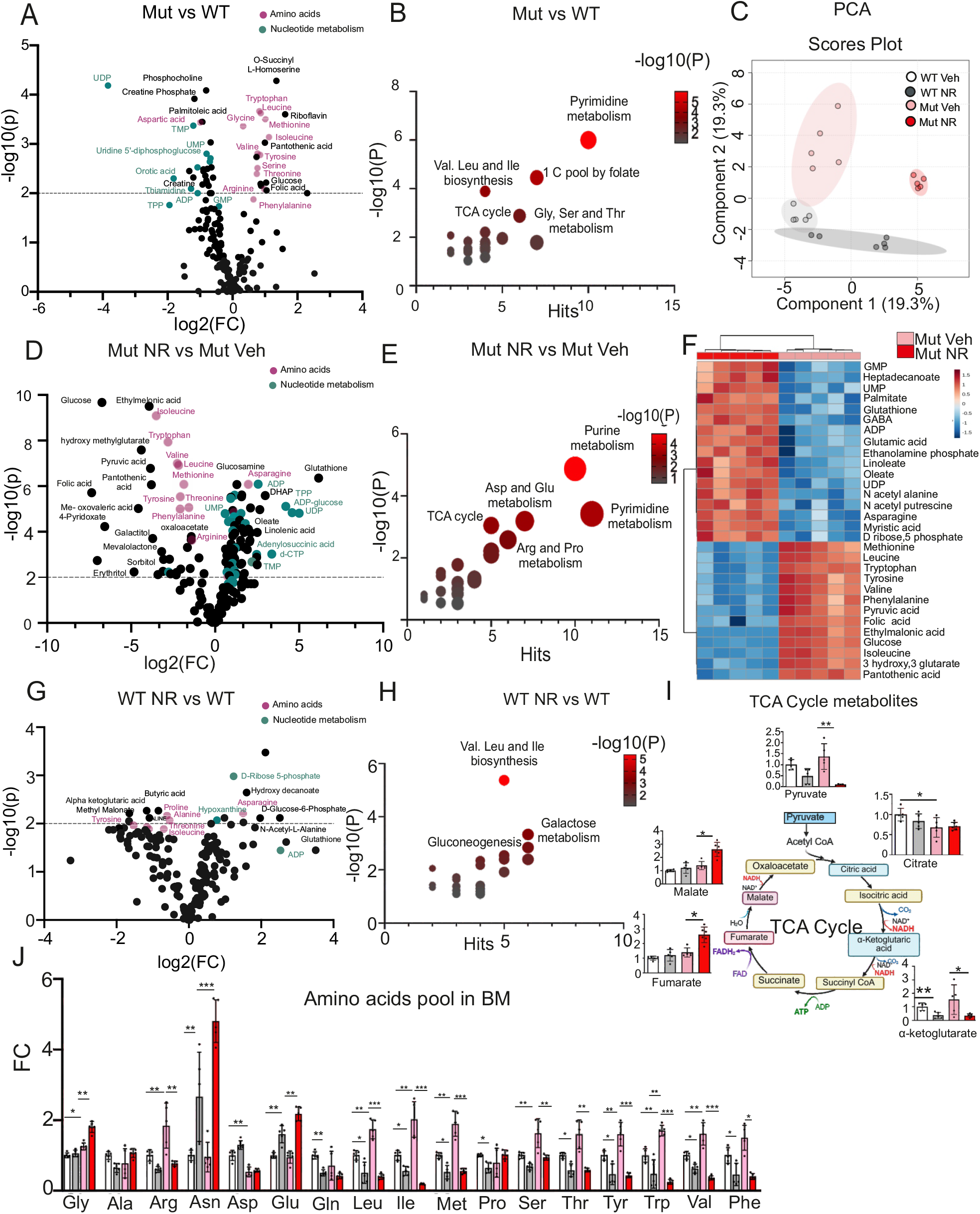
NAD^+^ treatment exacerbates anemia and impairs erythroid maturation in progeria mice. (A) NAD^+^ levels in blood of mutator mice (n = 4 mice per group). (B) NAD^+^ levels in blood following nicotinamide riboside (NR) supplementation (n = 6 mice per group). (C) Hemoglobin levels in mice over the course of treatment (x-axis indicates weeks) (n = 10 mice per group). (D-G) Hemoglobin (Hb) levels, red blood cell count (RBC), hematocrit (HCT) and white blood percentage by terminal blood analysis showing (WBC) (n = 7 mice per group). (H) Percentage of Thiazole Orange-positive cells in peripheral blood by flow cytometry analysis (n = 6 mice per group). (I) Percentage of Mitotracker Green-positive cells in peripheral blood by flow cytometry analysis (n = 6 mice per group). (J) Total TMRM intensity in reticulocytes in peripheral blood by flow cytometry analysis (n = 6 mice per group). (K, L) Active mitochondria based on Mitotracker green and TMRM positive cells in peripheral blood (n = 6 mice per group). (M-O) NAD^+^, NADH levels and NAD^+^/NADH ratio in harvested bone marrow cells (WT Veh: n = 5, WT NR: n = 5, Mut Veh: n = 6, Mut NR: n = 6). (P) Histological analysis of iron (blue) staining in the spleen red pulp to assess iron accumulation. Scale bars represent 100 µm. (Q-S) NADP^+^, NADPH levels and NADPH/NADP^+^ ratio in bone marrow cells (WT Veh: n = 5, WT NR: n = 5, Mut Veh: n = 6, Mut NR: n = 6). (T) Heatmap of NAD/P biosynthesis genes expression in BM of mutator and WT mice with Veh and NR treatment (WT Veh: n = 4, WT NR: n = 5, Mut Veh: n = 7, Mut NR: n = 7) Data are presented as mean ± SEM. Statistical analyses were performed using Student’s t-test or two-way ANOVA as indicated. ^*^p ≤ 0.05, ^**^p ≤ 0.01, ^***^p ≤ 0.001, ^****^p ≤ 0.0001.

### NR treatment elevates circulating immature erythrocytes and affects NADH pool in bone marrow

Mutator mice have been shown to exhibit circulating immature erythrocytes containing mitochondria and ribosomes in their blood consistent with delayed erythroid maturation^35^.To investigate whether NR treatment exacerbates this defect and alters the number of circulating immature erythrocytes, we collected peripheral blood samples from tail vein of mice after 21 weeks of NR treatment. NR-treated mutator mice exhibited an increase in immature erythrocytes, as indicated by elevated ribosome content by thiazole orange staining (Fig. 1H) and increased mitochondrial amount as determined by Mitotracker green staining (Fig. 1I). Furthermore, NR-treated reticulocytes showed heightened mitochondrial membrane potential, as evidenced by increased TMRM fluorescence (Fig. 1J–L), indicative of respiration competent mitochondria, a hallmark of incomplete erythrocyte maturation. This accumulation of immature erythrocytes coincided with enhanced erythrocyte clearance and degradation, as demonstrated by iron accumulation in the spleen, which was further elevated in NR-treated mutator mice compared to untreated counterparts (Fig. 1P). Conversely, bone marrow iron stores were markedly depleted in mutator mice (Fig. S1I), potentially limiting the availability of iron for heme biosynthesis and effective erythropoiesis. Together, these findings indicate that long-term NAD^+^ supplementation in the context of mitochondrial progeria exacerbates defective erythroid maturation, promoting premature erythrocyte clearance and splenic iron overload, key contributors to worsening anemia.

Redox metabolite profiling in bone marrow cells revealed a significant shift in the NAD^+^/NADH balance in mutator mice. Compared to controls, mutator mice exhibited elevated NADH levels in bone marrow cells (Fig. 1M–N). NR-treated mutator mice showed a disproportionate accumulation of NADH, resulting in a markedly reduced NAD^+^/NADH ratio (Fig. 1O). Furthermore, NR treatment decreased NADP^+^ in both genotypes (Fig. 1Q) but elevated NADPH levels in mutators (Fig.1 R,S) indicating a shift toward a more reductive state. Gene expression data from bone marrow shows decrease in Nicotinamide mononucleotide adenylyltransferase 3 (*nmnat3)* and *slc25a51* in NR treated mutator, while increase in nicotinamide phosphoribosyltransferase *(nampt)* (Fig 1T). To investigate whether these redox changes affected mitochondrial respiratory chain function, we assessed oxygen consumption of bone marrow cells with different substrates for respiratory chain complexes, in coupled and uncoupled state. Mutator mice exhibited reduced ADP-dependent respiration, lowered uncoupled respiration and slightly reduced inhibition of respiration with rotenone, a complex I inhibitor, suggesting mild CI-dependent respiration deficiency (Fig S1 G, H). NR treatment did not induce significant changes in CI-dependent respiration, although bone marrow cells from both wild-type and mutator mice treated with NR displayed a slight increase in overall respiration. Therefore, the mere deficiency in NADH/NAD+ recycling at CI is not likely to explain the NADH increase.

Mutator mice have been shown to have an increased number of erythroid progenitors and decreased B cell progenitors in their bone marrow^35^.We asked whether NR-mediated redox shift in bone marrow environment affects hematopoietic progenitor differentiation in control or mutator mice. Bone marrow harvested after 6 months of NR treatment showed no significant change in mutator erythroid progenitor phenotype (Fig S2A) or significant effect on B or T cell phenotype (Fig S2C). No change was observed in mitochondrial amount as seen by MTG staining in erythroid and myeloid lineage populations (Fig S2B and D). Although no change in cellularity of bone marrow was observed (Fig S2E) mutator mice displayed reduced hematopoietic stem cell (HSC) quantity (Fig S2F) which remained unchanged due to NR, furthermore no differences were observed in active mitochondrial potential within HSCs, as assessed by TMRM staining (Fig. S2G-H).NR treatment did not further alter long-term or short-term HSC populations (Fig S2I,J). Together, these findings indicate that while NR supplementation profoundly alters the redox landscape of the bone marrow marked by elevated NADH and NADPH levels, it does not affect mitochondrial mass or hematopoietic lineage distribution. Instead, the adverse hematologic effects of NR appear to stem from metabolic and reductive stress, rather than shifts in progenitor composition or mitochondrial content.

### Redox imbalance shifts the mutator bone marrow metabolism with glucose and folate depletion

To evaluate the metabolic consequences of redox imbalance, we performed targeted metabolomic profiling of 180 metabolites in the bone marrow cells from control and mutator mice. The changes in mutator mice include: 1) elevated steady-state levels of amino acids except for Ala, Asn, Glu, Gln, Pro (Fig. 2A, B, J); 2) decrease in nucleotide sugars and sugar phosphates with especial decrease of phosphorylated metabolites (Fig 2A, B); 3) creatine and phosphocreatine deficiency. These findings suggest a profound deficiency of cellular high-energy phosphates, affecting nucleotide synthesis, energy metabolism and stalling protein synthesis in the proliferative bone marrow of the untreated mutator mice.

**Figure 2.**
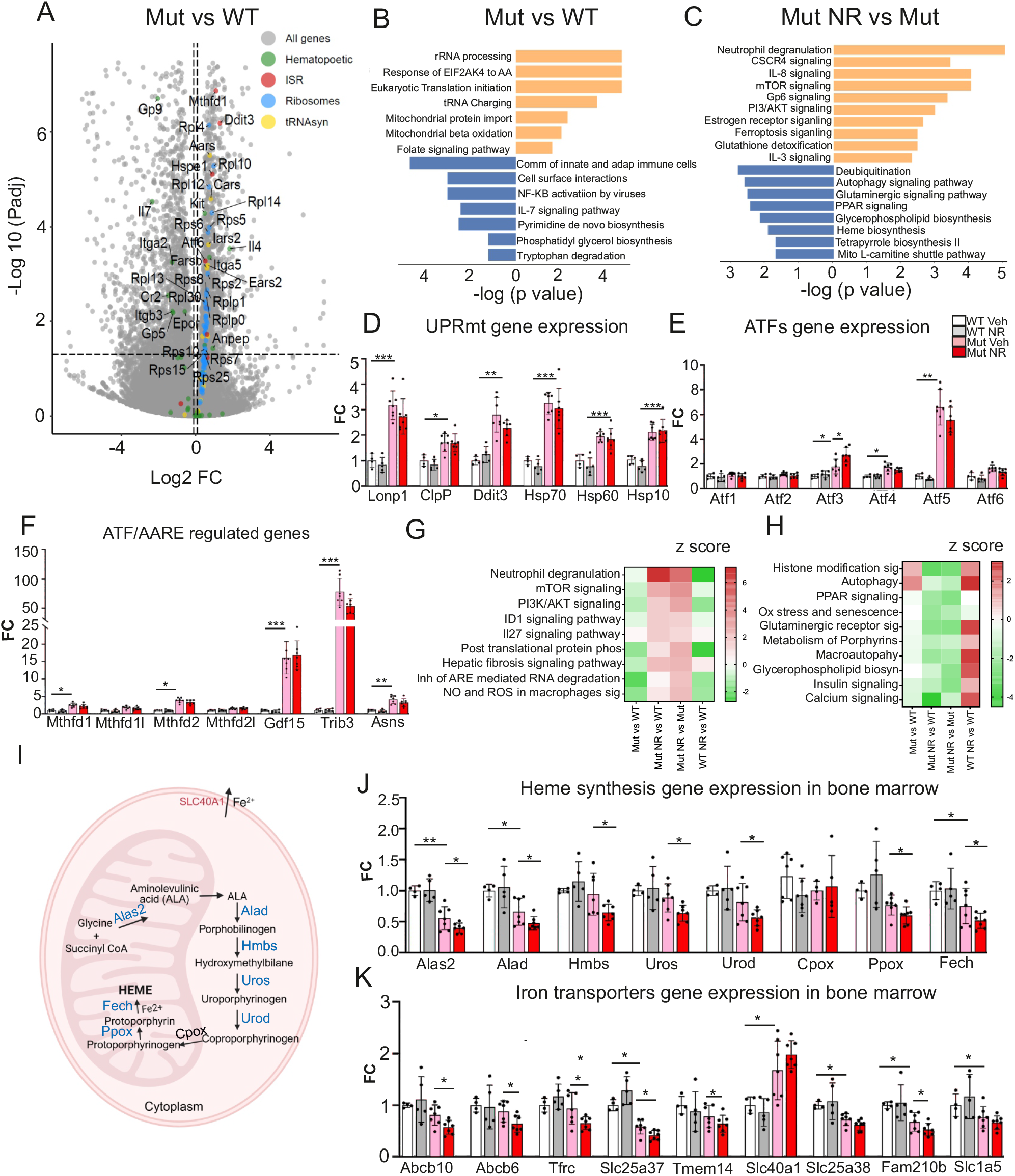
NR remodels metabolic profile in proliferative bone marrow cells. (A) Volcano plot depicting significantly altered metabolites in mutator bone marrow compared to control (wt) mice (n = 5 mice per group). (B) Significantly changed metabolite pathway in the bone marrow of mutator mice compared to control (n = 5 mice per group). (C) Principal Component Analysis (PCA) of bone marrow of bone marrow metabolites in mutator and control mice, treated with vehicle or nicotinamide riboside (NR) (n = 5/6 mice per group). (D) Volcano plot depicting significantly changed metabolites in NR-treated mutator bone marrow cells compared to vehicle-treated mutators (n = 5 mice per group). (E) Significantly changed metabolite pathway for the top metabolites in bone marrow of mutator mice, NR treatment and vehicle (n = 5 mice per group). (F) Heat map of the top 30 metabolites in mutator bone marrow, comparing vehicle and NR treatments (n = 5 mice per group). (G) Volcano plot showing significantly changed metabolites in NR-treated mutator bone marrow compared to vehicle-treated controls (n = 5 mice per group). (H) Significantly changed metabolite pathway for the top metabolites in bone marrow of NR-treated control mice compared to vehicle-treated controls (n = 5 mice per group). (I) TCA cycle metabolite levels in wt and mutator mice with vehicle or NR treatment (n = 5/6 mice per group). (J) Levels of amino acids in bone marrow, determined by targeted metabolomics, with fold-change normalized to wt (n = 5/6 mice per group). Data are presented as mean ± SEM. Statistical significance was assessed using Student’s t-test or two-way ANOVA, as indicated; ^*^p ≤ 0.05, ^**^p ≤ 0.01, ^***^p ≤ 0.001, ^****^p ≤ 0.0001.

The highly abnormal metabolic profile of mutator bone marrow prompted us to test the effects of NR treatment. Principal component analysis (PCA) revealed a distinct clustering of NR-treated mutator samples, separate from untreated mutators and controls (Fig. 2C), reflecting extensive metabolic reprogramming. Most prominent metabolic changes in NR treated mutator mice included (Fig 2D-F,J,): 1) Most amino acids were decreased to levels comparable to or significantly lower than those in wild-type controls, with the notable exception of aspartate and glutamine, the key substrates of asparagine synthetase (ASNS), which catalyzes the synthesis of asparagine and glutamate. Both asparagine and glutamate were significantly elevated following NR treatment in mutator mice, and to a lesser extent in wild-type mice. The levels of glycogenic branched-chain amino acids were remarkably decreased. Notably, tryptophan, a precursor in the de novo NAD^+^ biosynthesis pathway, was also reduced in NR-treated mutator bone marrow cells (Fig. 2J). Consistent with this, key intermediates of the tryptophan– kynurenine–NAD^+^ pathway, including kynurenine and quinolinic acid, were similarly decreased (Fig. 2D, J), suggesting either feedback suppression or reduced metabolic flux through the de novo NAD^+^ biosynthesis pathway under chronic NR supplementation; 2) elevation of nucleotide sugars and sugar phosphates; 3) depletion of steady-state glucose and pyruvate suggesting increased usage for glycolytic metabolism; 4) partially stalled TCA cycle, inhibiting oxidative energy metabolism (Fig 2I). High NADH inhibits malate dehydrogenase leading to increase of fumarate and malate; 5) High glutathione suggests increased reductive power and 6) accumulation of lipids (palmitic acid, oleic acid, linolenic acid, and myristic acid; Fig 2D,F) suggest stalled β-oxidation as oxidative metabolism is inhibited. Depletion of folate has the potential to severely dampen cellular biosynthesis reactions from translation to lipid and nucleotide synthesis in rapidly dividing cells. Treated mutator showed stronger shift in metabolic profile away from untreated control (Fig S3A,B). NR-treated wild type mice showed mildly decreased amino acid levels, except for a marked increase in asparagine and glutamate, along with elevated levels of sugar phosphates (Fig. 2G, H). However, there was no corresponding increase in nucleotide biosynthesis or lipid accumulation. Taken together, these findings suggest that NR treatment, and the resulting accumulation of NADH, imposes a metabolic burden on bone marrow cells by severely depleting glucose, possibly due to its diversion into glutathione synthesis via the serine/glycine pathway, or due to reduced glucose uptake. In addition, oxidative glucose metabolism is impaired by a partially stalled TCA cycle along with depletion of folate, the central one-carbon carrier essential for biosynthesis.

### NR enhances inflammation in mutators without affecting metabolic stress responses

We next examined gene regulatory pathways altered in the mutator bone marrow. Notably, genes encoding components of the translation machinery including aminoacyl-tRNA synthetases, ribosomal subunits, and ribosome assembly factors were significantly upregulated (Fig. 3A, B; Fig. S3C, E). The upregulation of cytoplasmic ribosomal genes was more pronounced than that of mitochondrial ribosomal genes, suggesting a potential uncoupling of nuclear and mitochondrial translation programs (Fig. S3C). In parallel, mitochondrial stress responses were activated, including both the mitochondrial integrated stress response (ISR^mt^) and the unfolded protein response (UPR^mt^) (Fig 3D, F). Common upstream regulators of these pathways, the ATF transcription factors, were induced: ATF5 showed a 6-fold increase, ATF4 a 1.5-fold increase, and ATF3 a 2-fold increase (Fig. 3E). Correspondingly, ATF-regulated ISRmt target genes were strongly upregulated, including *Gdf15, Trib3, Asns, Mthfd1, and Mthfd2*, implicating coordinated activation of amino acid and redox stress signaling (Fig. 3F). Further evidence of disturbed proteostasis was provided by the upregulation of UPR^mt^ components, including *LonP, ClpP*, and several mitochondrial chaperones (Fig. 3D). Importantly, NR treatment did not suppress any of these stress responses (Fig. 3D–F); in fact, it further elevated Atf3 expression in NR-treated mutator mice (Fig. 3E), consistent with ATF3’s role as a redox- and nutrient-sensitive transcription factor often associated with late-stage cellular stress or terminal differentiation.

**Fig 3.**
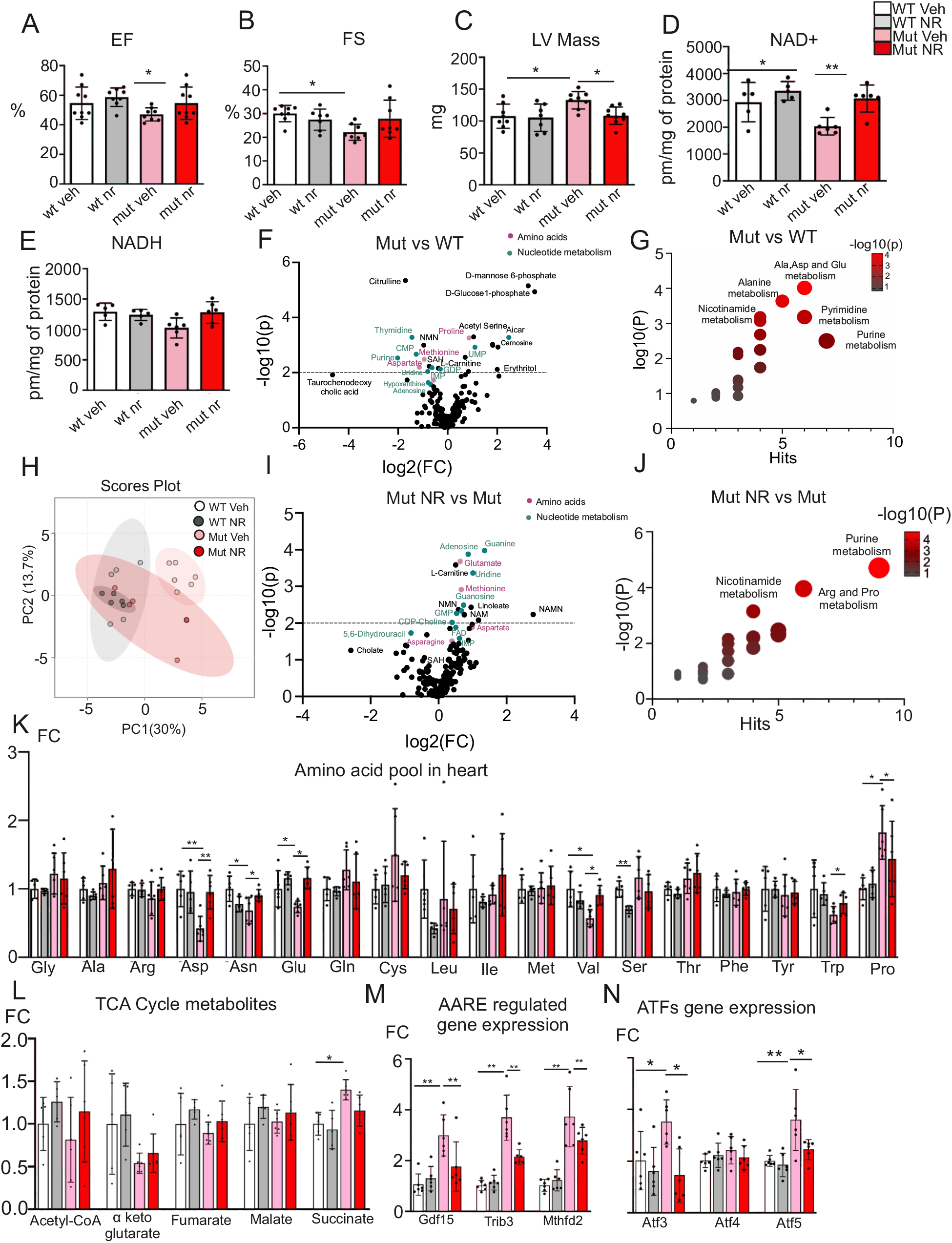
NR enhances inflammation and suppresses heme biosynthesis in mutator mice. (A) Transcriptome profile of bone marrow from mutator and wild-type (wt) mice, presented as a volcano plot showing significance (adjusted p-value, Padj) and fold change (FC). (B) Top pathways changed in the bone marrow transcriptome in NR-treated control mice compared to vehicle-treated control mice by ingenuity pathway analysis (WT Veh: n = 4, Mut Veh: n = 7) (C) Top pathways changed in the bone marrow of NR-treated mutator compared to vehicle-treated mutator mice by ingenuity pathway analysis (Mut Veh: n = 7, Mut Nr: n = 7) (D) Transcript levels of ISRmt genes in the bone marrow of wt and mutator mice with vehicle or NR treatment (WT Veh: n = 4, WT NR: n = 5, Mut Veh: n = 7, Mut NR: n = 7). (E) Transcript levels of ATF genes in the bone marrow of wt and mutator mice with vehicle or NR treatment (WT Veh: n = 4, WT NR: n = 5, Mut Veh: n = 7, Mut NR: n = 7). (F) Transcript levels of UPR^mt^ genes in the bone marrow of wt and mutator mice with vehicle or NR treatment (WT Veh: n = 4, WT NR: n = 5, Mut Veh: n = 7, Mut NR: n = 7). (G,H) Heatmap showing z-scores for selected canonical pathways. Z-scores were calculated by IPA to predict pathway activation (red) or inhibition (green) based on gene expression changes. (I) Schematic representation of heme biosynthesis created using Biorender. (J) Transcript levels of heme biosynthesis genes in the bone marrow of wt and mutator mice with vehicle or NR treatment (WT Veh: n = 4, WT NR: n = 5, Mut Veh: n = 7, Mut NR: n = 7). (K) Transcript levels of heme and iron transporter genes in the bone marrow of wt and mutator mice with vehicle or NR treatment (WT Veh: n = 4, WT NR: n = 5, Mut Veh: n = 7, Mut NR: n = 7). Data represent results from IPA pathway analysis using input gene sets from each comparison. Data are shown as mean ± SEM. Statistical significance was determined using Student’s t-test or two-way ANOVA where indicated; ^*^p ≤ 0.05, ^**^p ≤ 0.01, ^***^p ≤ 0.001, ^****^p ≤ 0.0001.

The gene signatures after NR treatment of mutators were especially emphasizing immune activation, including upregulation of pathways involved in neutrophil degranulation, *Cxcr4*, IL-8, and IL-3 signaling (Fig. 3C), whereas these responses were absent in NR-treated wild-type mice (Fig. S3D), indicating a disease- and tissue-specific inflammatory effect of NR supplementation. In addition, NR-treated mutator mice showed significant upregulation of mTOR signaling, a central regulator of anabolic metabolism, in both mutator and wild-type bone marrow (Fig. 3C, S3D). Pathway enrichment analysis further revealed that NR-treated mutators, compared to both vehicle-treated mutators and control mice, exhibited increase of mTOR pathways and downregulation of autophagy, PPAR signaling, insulin signaling, calcium signaling, and porphyrin metabolism (Fig. 3G, H). These results demonstrate that long-term NAD^+^ supplementation in mutator mice triggers disease-specific transcriptional changes, characterized by enhanced anabolic signaling, immune activation, and dysregulation of stress response coordination in proliferative bone marrow cells.

### NR impairs heme biosynthesis and disrupts iron handling in bone marrow

Given the observed reduction in hemoglobin (Hb) levels in NR treated mutator mice, we next investigated the impact of NAD^+^ supplementation on the heme biosynthesis pathway. Mutator mice exhibited a significant downregulation of 5′-aminolevulinate synthase 2 (*Alas2*), the mitochondrial enzyme catalyzing the rate-limiting first step in porphyrin ring synthesis (Fig. 3I,J). Expression of multiple enzymes involved in the downstream steps of heme biosynthesis was also markedly reduced in mutator mice and was further suppressed by NR treatment, indicating a profound disruption of heme metabolism. In contrast, wild-type mice showed no significant changes in heme biosynthesis gene expression following NR supplementation (Fig. 3I,J). This impairment in heme synthesis was accompanied by dysregulation of mitochondrial iron transport. Expression of *Abcb6, Abcb10*, and *Slc25a37* (Mitoferrin-1) genes critical for mitochondrial iron import and porphyrin transport was significantly reduced in mutator mice and further downregulated upon NR treatment (Fig. 3K). Notably, *Abcb10* stabilizes Mitoferrin-1 and facilitates ferrous iron import into mitochondria, a prerequisite for both heme and iron–sulfur (Fe–S) cluster synthesis. Similarly, *Slc25a38*, a mitochondrial glycine transporter essential for the first committed step of heme biosynthesis, and *Tfrc*, encoding the transferrin receptor that mediates cellular iron uptake, were both significantly reduced in expression in mutator mice and further suppressed by NR (Fig. 3K). Interestingly, *Slc40a1*, encoding the iron exporter ferroportin, was significantly upregulated in mutator mice, with a modest further increase following NR treatment. Collectively, these data indicate that mitochondrial dysfunction in mutator mice disrupts both heme biosynthesis and iron homeostasis, critical processes for effective erythropoiesis. Importantly, chronic NAD^+^ supplementation exacerbates these impairments, further compromising red blood cell development and contributing to anemia in the context of mitochondrial disease.

### NR treatment improves cardiac NAD^+^ levels and heart function in mutator mice

Mutator mice develop age-dependent cardiomyopathy by 1 year of age^36,37^, our echocardiographic analysis evidenced their reduced ejection fraction (EF) decreased fractional shortening (FS) and an increase in left ventricle (LV) mass (Fig. 4A-C). Interestingly, NR treatment significantly improved these cardiac functions in the mutator mice: LV mass was reduced and EF and FS improved (Fig. 4A-C). These findings indicate that NR treatment mitigates and delayed the progression of cardiomyopathy.

**Figure 4.**
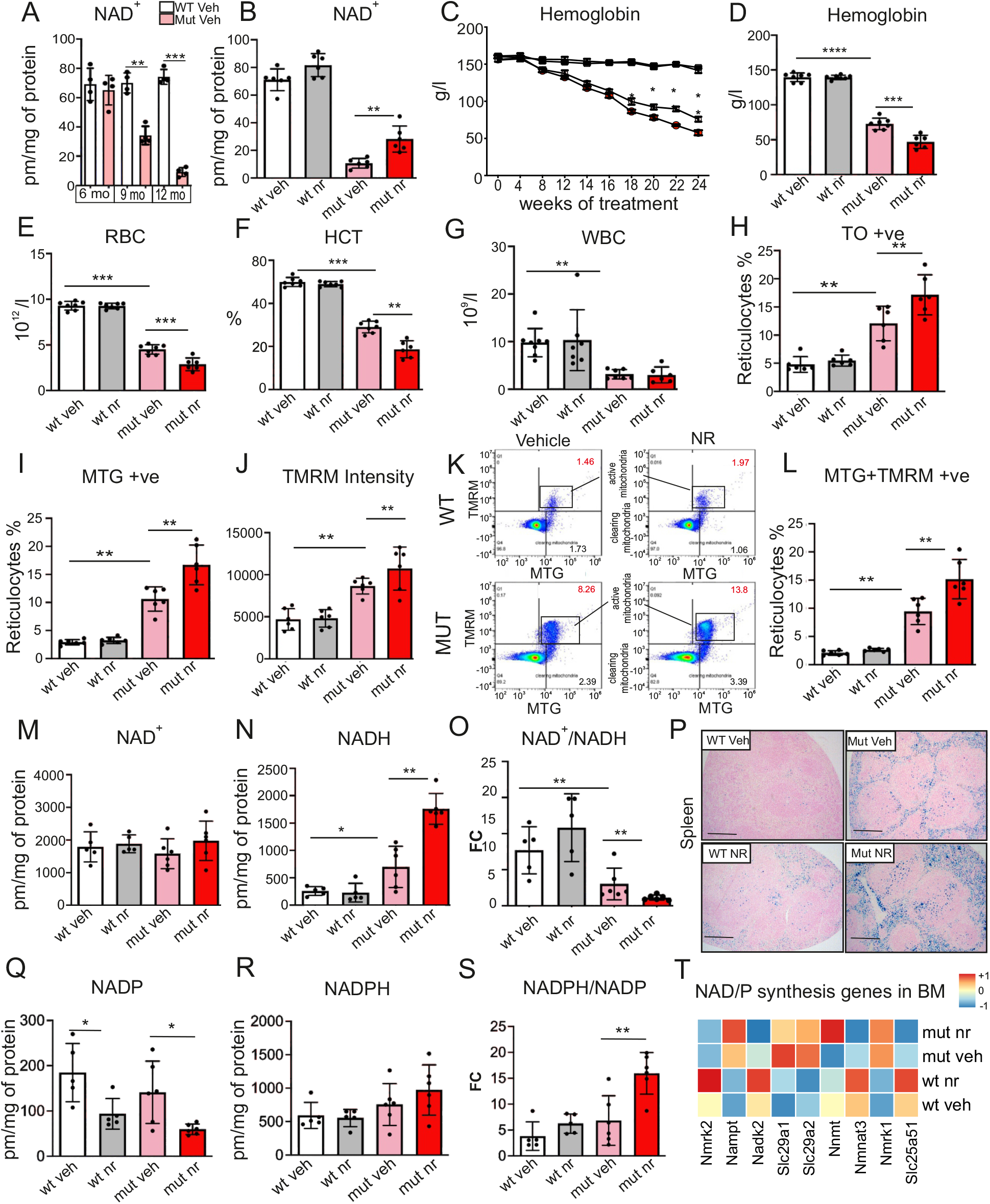
NR treatment improves cardiac function and metabolic profile in mutator mice. (A-C) Cardiac function assessed by echocardiography with vehicle and NR treated wild-type and mutator mice showing (A) ejection fraction (EF), (B) fractional shortening (FS), and (C) left ventricular mass (LV mass). (n = 8 mice per group). (D-E) NAD+ and NADH levels in heart tissue of mutator and WT mice treated with vehicle or NR (n = 6 mice per group). (F) Volcano plot showing significantly changed metabolites in the heart of mutator mice compared to WT mice (WT Veh: n = 5, Mut Veh: n = 6). (G) Significantly changed metabolite pathway for the top metabolites in the heart of mutator mice (WT Veh: n = 5, Mut Veh: n = 6). (H) Principal Component Analysis (PCA) of metabolomic data from the heart of mutator and WT mice treated with vehicle or NR (WT Veh: n = 5, WT NR: n = 5, Mut Veh: n = 6, Mut NR: n = 6). (I) Volcano plot showing significantly changed metabolites in NR-treated mutator heart compared to vehicle-treated mutators (n = 6 mice per group). (J) Significantly changed metabolite pathway for the top metabolites in NR-treated mutator heart (n = 6 mice per group). (K) Amino acid levels in heart WT and mutator mice with vehicle or NR treatment assessed by targeted metabolomics, with fold-change normalized to WT (WT Veh: n = 5, WT NR: n = 5, Mut Veh: n = 6, Mut NR: n = 6). (L) TCA cycle metabolites in WT and mutator mice with vehicle or NR treatment (WT Veh: n = 5, WT NR: n = 5, Mut Veh: n = 6, Mut NR: n = 6). (M-N) Transcript levels of ISR^mt^ and ATF genes in the heart tissue of wt and mutator mice with vehicle or NR treatment (n= 6 mice per group). Data are presented as mean ± SEM. Statistical significance was assessed using Student’s t-test or two-way ANOVA where indicated; ^*^p ≤ 0.05, ^**^p ≤ 0.01, ^***^p ≤ 0.001, ^****^p ≤ 0.0001.

Analysis of cardiac NAD^+^ metabolism revealed that mutator hearts had reduced NAD^+^ level at baseline, which was restored by NR treatment (Fig. 4D). NADH levels remained stable in mutator hearts (Fig. 4E), maintaining a normal NAD^+^/NADH ratio (Fig. S4A), in stark contrast to the redox imbalance in the bone marrow. Both NADP^+^ and NADPH levels were mildly decreased (Fig. S4B–D), but glutathione levels remained unchanged (Fig. S4E–G), indicating preserved redox homeostasis in the heart. Targeted metabolomic profiling following the same method as for bone marrow revealed that NR treatment led to broad normalization of cardiac metabolic signatures in mutator mice. Metabolites involved in amino acid metabolism, purine biosynthesis, and redox pathways were significantly altered in untreated mutator hearts and partially restored toward control levels following NR supplementation (Fig. 4F–J). Principal component analysis (PCA) confirmed a metabolic shift in NR-treated mutator hearts toward the control like profile, contrasting with the divergent remodeling observed in bone marrow (Fig. 4H). NR-treated mutator mice showed an increase in the steady-state levels of NAD^+^ biosynthesis intermediates (NAM, nicotinamide; NMN, nicotinamide mononucleotide; NAMN, nicotinate mononucleotide), which are part of the salvage and Preiss-Handler pathways of NAD+ metabolism (Fig 4I, J). The steady state levels of amino acids and TCA cycle metabolites mostly were similar to controls, but proline remained high in the untreated and treated mutators (Fig 4K, L). NR also restored the steady-state level of L-carnitine, a critical for cardiac energy metabolism, particularly fatty acid oxidation. At the transcriptional level, mutator hearts induced ISR^mt^ signature (*Gdf15, Trib3, Mthfd2, Atf3, Atf4*, and *Atf5*) under basal conditions, which was resolved by NR treatment (Fig. 4M–N), further pointing to restoration of both cardiac function and metabolism.

To further evaluate the physiological benefits of NR, we assessed the motor coordination and systemic metabolism of mutators and their controls. Both mutator and wild type mice showed a trend towards improved motor performance in rotarod test after NR-treatment compared to controls (Fig. S4L). Metabolic cage analysis showed elevated oxygen consumption, increased CO_2_ production, and a modestly higher respiratory exchange ratio (RER) in NR treated mutator and control mice indicating a shift toward glucose utilization (Fig. S4M-O) which was not affected by NR-treatment. Together, these results show that long term NR supplementation slows down development of cardiomyopathy in mutator mice and restores metabolism, whilst being deleterious for the bone marrow.

## Discussion

NAD^+^ and its derivatives have a central role in energy metabolism, biosynthesis pathways and physiological responses. Therefore, the reports of NAD^+^ depletion in different mitochondrial, metabolic, and age-associated disorders ^12,38,39^ and the rescuable nature of the deficiency have raised high academic and commercial interests in NAD^+^ booster treatments. Currently, over 100 registered clinical trials (ClinicalTrials.gov) are ongoing using NAD^+^ boosters in both rare and common diseases. Still, insufficient information exists of tissue-specific responses to long-term increase of NAD^+^ augmentation. Here, we studied the effects of NAD-booster nicotinamide riboside in mitochondrial progeria mice that show NAD^+^ depletion and disease phenotypes in both solid and proliferative tissue types. We report that NAD booster nicotinamide riboside treatment improves cardiac function and prevents the development of cardiomyopathy in mitochondrial progeria mice but worsens their anemia. NR has dual effects in solid and proliferating tissue types: NAD^+^ content increases in the heart, but in the bone marrow, it causes accumulation of reduced form of NAD metabolites, NADH and NADPH. The latter causes reductive stress and aggravates erythroid differentiation defect of hematopoietic stem cells, leading to rapidly progressing anemia. The findings underscore the importance of knowledge on how strong metabolic modifiers such as NR affect tissues with different metabolic needs. Our study was prompted by previous findings in mitochondrial myopathy patients with systemic NAD^+^ depletion^16^. Long-term niacin therapy improved muscle strength but caused hemoglobin to decrease in patients ^16^. Similarly, short-term high-dose NR supplementation reduced hemoglobin, hematocrit, and platelet counts in healthy individuals^40^ pointing indirectly to NAD^+^ booster effects on erythroid lineage. Our current findings underscore the importance of NAD^+^ dependent redox homeostasis in erythrocyte maturation and how it is modified by NAD^+^ precursor treatment.

Despite the partial restoration of the systemic NAD^+^ deficiency in blood of mutator mice, NR aggravates erythroid maturation defect, resulting in premature release of immature erythrocytes in the circulation. These become subsequently sequestered and degraded in the spleen, with redistribution and imbalance of iron pool^35^, depleted in bone marrow and accumulated in the spleen. Bone marrow is a highly dynamic and proliferative tissue, characterized by rapid cell turnover, making it particularly sensitive to changes in metabolism and nutrient availability^41,42^. Previous studies have shown that proliferative cells have a demand for intracellular oxidized NAD^+ 43–45^ making them more vulnerable to redox changes. We observed a marked redox imbalance in NR-treated mutator bone marrow, characterized by NADH and NADPH accumulation. The redox shift towards a lower NAD^+^/NADH and higher NADPH/NADP^+^ ratios disrupt metabolic homeostasis and alters the balance between anabolic and catabolic processes in cells^11,46,47^. This is evident by increase in metabolites associated with purine and pyrimidine metabolism, alongside a significant depletion of glucose and pyruvate, suggesting a shift towards glycolytic metabolism. The bone marrow of mutator mice also showed lowered folate levels, the major carrier of glucose-derived one carbon units, despite normal folate intake. This finding is shared with patients on long term niacin who also showed lower serum folate levels^16^. Overall, the data highlight the critical role of a tightly regulated redox state, and interplay between B-vitamin derived cofactors, including NAD^+^ precursor B3 and folate B9 in maintaining metabolic homeostasis, especially in tissues with high cellular turnover.

The hypertrophic cardiomyopathy in the mutator mice was associated with lowered levels of NAD^+^ metabolites, NAM, NMN and NAMN, the depletion of which was prevented by NR treatment. Indeed, NR supplementation was able to sustain cardiac contractile function in aging mutator mice. These findings align with recent studies on NAD^+^ booster effects in the heart in different model systems, enhancing oxidative metabolism, mitochondrial biogenesis and cardiac function^14,21,23,57^. The elevated levels of L-carnitine in NR-treated mutator hearts support the conclusion of improved mitochondrial oxidative capacity and beta-oxidation. Furthermore, NR prevented induction of ISR^mt^ that was upregulated in the untreated mice, indicating that correction of NAD^+^ metabolism by NR delayed or prevented manifestation of cardiomyopathy in the mice whose lifespan was, however, limited by the anemia aggravated by NR. In the bone marrow, however, the mitochondrial integrated stress response^51,54,55^ and unfolded protein response (UPRmt)^52,53,58^,were induced similarly in non-treated and NR-treated mice, indicating that both responses are markers of disease severity and not redox-regulated in a pathological context.

While our study was conducted in a progeric mouse model, the anemia observed in these mice closely resembles that seen in Pearson syndrome, a severe, childhood-onset anemic disorder caused by mtDNA deletions. In Pearson syndrome, in both mice and patients, immature erythrocytes are released in circulation^35^. Anemia is also reported to be an important component of many mitochondrial diseases and previously reported to also associate with poor prognosis^59^.

The evidence emphasizes the importance of close follow up of hematologic parameters when treating patients with NAD+ boosters, particularly in those with mitochondrial disease. Future work on refining NAD^+^ repletion strategies is needed, to avoid deleterious effects in vulnerable tissues. This includes optimizing dosing regimens and exploring combinatorial approaches to buffer redox shifts or support biosynthetic pathways. As NAD^+^ boosters enter late-stage clinical trials and potential approval, a mechanistic framework that incorporates tissue-specific vulnerabilities and benefits will be critical to guide their safe and effective use.

### Limitations of the study

While this study provides evidence for tissue-specific responses to NAD^+^ restoration under mitochondrial stress, the mitochondrial mutator mouse is a complex model with energy metabolic and stem cell homeostatic defect, leading to premature aging. As such, the extent and nature of redox and metabolic imbalances observed may not fully capture the spectrum of responses in more moderate or chronic disease settings. Future studies involving patients, especially those receiving long-term NAD^+^ supplementation, will be essential to validate these findings and to define the therapeutic window, safety, and physiological relevance of NAD^+^ repletion across tissues. Nonetheless, our results establish a crucial mechanistic framework that underscores the need for context-dependent NAD^+^ interventions and highlight the risks of assuming uniform benefit across diverse tissue and disease environments.

## Research Methodology

Resource availability

## Lead contact

Further information and requests for resources and reagents should be directed to and will be fulfilled by the lead contact

## Materials availability

This study did not generate any new material

## Data and code availability

The Gene expression data generated for this study have been deposited in the European Nucleotide Archive (ENA) under the accession number PRJEB90288. The dataset is publicly available and can be accessed at: https://www.ebi.ac.uk/ena/browser/view/PRJEB90288.

The metabolomics data from this study have been deposited in the MassIVE repository under the accession number MSV000098316. The dataset can be accessed at: https://massive.ucsd.edu/ProteoSAFe/dataset.jsp?accession MSV000098316. This paper does not report original code.

Additional information required to reanalyze the data reported in this paper is available from the lead contact upon request.

## Experimental models

### Mouse models

The Ethical Review Board of Finland approved all animal experimentation. Mutator mice for this study is a knock-in inactivating mutation (D257A) in the exonuclease domain of DNA polymerase gamma were used ^33^.The strain was maintained by crossings with *D257A/wt* males and *wt/wt* (B57BL/6JRcc) females to prevent transmission of mtDNA mutations to germline. Experimental groups were produced by crossings with F1 generation *D257A/wt* females and *D257A/wt* males. Five months old mice were given nicotinamide riboside (NR) provided by Chromadex, baked in powdered food for 24 weeks, Vehicle treated mice were given chow diet from same commercial provider without NR as in previous study by^17^. Each mice received the daily dose of 400mg/kg/day of NR during this study. Food intake in the treatment study was controlled by removing the food from cages 4 hr before the mice were sacrificed for sample collection.

### Hemoglobin measurement

Hemoglobin levels were monitored longitudinally in mice via peripheral blood sampling. A small drop of blood (∼5–10 µL) was collected from the tail vein every two weeks using a sterile scalpel or 27G needle following brief warming of the tail to promote vasodilation. The first drop was wiped to prevent contamination, and the subsequent drop was used for analysis. Hemoglobin concentration was measured immediately using a portable hemoglobinometer (Hemo Control-EKF diagnostics) with hemoglobin microcuvettes (EKF diagnostics), following the manufacturer’s instructions. All measurements were performed in duplicate to ensure accuracy, and mice were handled gently to minimize stress.

### Respiratory Function Testing in Animals

Oxygen consumption and carbon dioxide production were measured in Oxymax Lab Animal Monitoring System (CLAMS; Columbus Instruments). In CLAMS the mice were housed in individual cages in temperature-regulated chamber, settled for 24 hours and were recorded for 24 hours. The settling and the first day of recording were in room temperature (+22°C). Respiratory exchange ratio (RER) was calculated to indicate a rough estimate of the preferred fuel (carbohydrate breakdown vs. fatty acid oxidation) of the organism.

### Mouse Phenotyping and Grip Strength

Body weight was recorded throughout the study using a precision electronic balance calibrated for small rodents. Grip strength was assessed using the BIO-GS3 apparatus (Bioseb, France). Mice were gently placed on the grid platform, ensuring all four limbs were engaged. A steady, horizontal pull was applied until the animal released its grip, and the peak force was recorded. For each animal, five consecutive measurements were performed, and the average value was calculated and normalized to body weight (g/g) to account for size differences. To minimize variability due to learning effects, all animals underwent grip strength training for three consecutive days prior to data collection.

### Terminal blood samples and blood count

At the study endpoint, ice were euthanized and blood was collected via heart puncture into EDTA tube (Greiner bio-one). Blood count was performed from whole blood samples during the same day using Advia 2120i analyzer (Siemens). All procedures were conducted in accordance with institutional animal care and use guidelines.

### Bone Marrow collection

Bone marrow was harvested from the femur and tibia by flushing the cavities with ice-cold Dulbecco’s Modified Eagle Medium (DMEM) using a 23-gauge needle under sterile conditions. The resulting cell suspension was passed through a 70 µm cell strainer to remove debris and clumps. Total cell counts were determined using an automated cell counter (Beckman Coulter). Cells were then aliquoted, centrifuged to remove residual media, and the resulting pellets were snap-frozen in liquid nitrogen for downstream molecular and metabolic analyses.

### Oroboros analysis from bone marrow

Oxygen consumption rates were determined in an Oroboros O2k as described before^60^. Briefly, bone marrow cells were harvested by rinsing of the femur with 1xPBS using a 26-gauge needle attached to a 1 mL syringe. After pelleting, 5×106 cells were resuspended in respiration medium containing 0.5 mM EGTA, 3 mM magnesium chloride, 60 mM K-lactobionate, 20 mM taurine, 10 mM potassium dihydrogen phosphate, 20 mM HEPES, 110 mM sucrose and 1 g/L fatty acid-free BSA (pH 7.1). After assessing routine respiration, the cells were permeabilized using digitonin, and supplemented with malate, pyruvate and glutamate. Ccomplex I and complex II-dependent respiration were determined by addition of saturating ADP levels followed by succinate. Oligomycin (10 nM) was added to determine leak respiration. This was followed by titration (0.5 μM steps) of carbonylcyanide p-trifluoromethoxyphenyl-hydrazone (FCCP) to assess maximal uncoupled respiration. Complex I respiration was measured by adding rotenone (0.5 μM) and residual by addition of antimycin A (2.5 μM). Complex IV activity was obtained by addition of ascorbate (2 mM) and N,N,N,N-Tetramethyl-p-phenylenediaminedihydrochloride (TMPD) (0.5 mM), followed by azide (10 mM).

### Tissue redox profiling

Quantitative analysis of NAD+, NADH, NADP+, NADPH, and reduced and oxidized glutathione in frozen tissue and blood samples was done as a collaboration in NADMED laboratory (Helsinki, Finland) using a proprietary technology (for further information see www.nadmed.com). Samples were shipped on dry ice and were stored at −80°C prior analysis. Frozen piece of tissue was placed into lysis solution pre-warmed to 50°C and homogenized in the tube with ceramic beads using automated Precellys homogenizer. Lysis solution is a water-based complex mixture of organic solvents, which force protein unfolding at temperatures above +45° C and release of all non-covalently bound metabolites into solution. Next, the tube with homogenate was placed on ice to force protein precipitation followed by centrifugation at 20000g for 10 min at +4°C to remove protein pellet and separate the extract. Frozen blood samples were thawed on ice-water bath in strictly controlled temperature conditions for 10-12 min and 100 µl of whole blood was injected into extraction buffer equilibrated to 80°C. Resulting homogenate was incubated for 1 min at 80° C followed by cooling step on ice-water bath for protein precipitation and centrifugation as for the tissue homogenates. Next, NAD+, NADH, NADP+, NADPH, GSH pool, and GSSG were measured individually from every sample extract using modified cyclic enzymatic reactions with colorimetric detection^61^. For normalization of the results for tissue samples, protein content in the sample was measured using the pellet obtained after centrifugation of the homogenate.

### Cardiac Echocardiography

Cardiac function was analyzed using a Vevo 2100 Ultrasound System (Visual Sonics, Toronto, ON, Canada). The animals were anesthetized with isoflurane for maximum of 15 min during the imaging (induction: 4.5% isoflurane, 450 ml/min air, maintenance: 1.5% isoflurane, 200 ml/min air, Baxter International, Deerfield, IL). Left ventricular mass, ejection fraction, fractional shortening, interventricular spectrum width (diastolic and systolic), left ventricular internal diameter (diastolic and systolic), left ventricular posterior wall width (diastolic and systolic) and left ventricular volume (diastolic and systolic) were determined from the parasternal short-axis (SAX) M-mode measurements. EF was calculated by the Vevo software using Teicholz formula.

## Flow cytometric analysis

### TO/MTG/TMRM flow cytometry for peripheral blood

Peripheral blood was collected into EDTA-coated tubes and diluted 1:500 in reticulocyte culture medium. Reticulocyte maturation was assessed by staining ribosomal RNA with thiazole orange, following established protocols. Intensity of ribosomal RNA staining was used to stage reticulocytes as immature (most intense thiazole orange (TO) signal), intermediate and mature according to standard protocol. Fluorescent signal was detected by Accuri flow cytometer (Becton Dickinson) and 30,000 cells per sample were analyzed.Samples were analyzed using FACSAria II SORP (BD Biosciences) flow cytometer.

### Extraction of hematopoietic cells

Bone marrow was collected from femur and tibia by flushing using 23 G needle. Cells were suspended in staining medium, Hank’s Balanced Salt Solution (HBSS) supplemented with 2 % bovine serum and filtered through a 40 µm nylon cell strainer. Cells were counted using automated cell counter. Samples were kept on ice and forwarded to analysis during the day of extraction.

### Flow cytometry analysis of hematopoietic populations

For analyzing hematopoietic stem and progenitor cell populations cells were stained using fluorophore-conjugated antibodies against lineage markers (CD2, CD3, CD5, CD8, B220, Ter119 and Gr1), c-kit, Sca1, CD48, CD150, CD105, CD34, CD16/32 and CD41. Samples were analyzed using FACSAria II SORP (BD Biosciences) flow cytometer.

### MTG/TMRM of hematopoietic populations

For analyzing mitochondrial mass and membrane potential in hematopoietic populations cells were first stained using fluorophore-conjugated antibodies against lineage markers (CD2, CD3, CD5, CD8, B220, Ter119 and Gr1), c-kit, Sca1, CD48, CD150, CD105, CD34, CD16/32 and CD41. Mitochondrial dyes MTG and TMRM were added together with Verapamil, an ABC efflux pump inhibitor, to ensure proper staining of the HSCs. Samples were analyzed using FACSAria II SORP (BD Biosciences) flow cytometer.

### Iron staining in bone marrow and spleen

Spleens from 11 month-old mice and femoral bones were collected and fixed in 10% neutral-buffered formalin following standard protocols. Bones were subsequently decalcified in 10% EDTA (pH 7.4) at room temperature before routine paraffin embedding. Tissue sections (3– 5 μm) were cut onto glass slides for histological analysis. Iron accumulation in the spleen sections and bone marrow cells was assessed using Perl’s Prussian Blue staining. Paraffin-embedded sections were deparaffinized in xylene, rehydrated through a descending ethanol series, and rinsed in distilled water. Slides were then incubated for 20 minutes in a freshly prepared solution of 5% potassium ferrocyanide in 10% hydrochloric acid. After thorough washing in water, nuclear counterstaining was performed using 0.1% nuclear fast red (Kernechtrot). Finally, sections were dehydrated through an ascending ethanol series, cleared in xylene, and cover slipped for imaging.

## Metabolomics

### Sample preparation of heart tissue samples

Metabolites were extracted from approximately 20 mg mouse heart tissue sample with 500 μL of cold extraction solvent (Acetonitrile: Methanol: Milli-Q Water; 40:40:20). Samples were homogenized with three cycles of 30 seconds at 5000 rpm with 60 seconds pause in between at +4 °C using a homogenizer (Precellys Evolution, Bertin Technolgies, France). The samples were then centrifuged at 14 000 rpm for 10 min at +4 °C, after which the supernatants (350 µl) were transferred to evaporation tubes and gently evaporated dry under nitrogen stream. Dried samples were resuspended in 40 µl cold extraction solvent (Acetonitrile: Methanol: Milli-Q Water; 40:40:20) and transferred to HPLC auto sampler glass vials.

### Sample preparation of bone marrow cells

Metabolites were extracted from 5 million cells, by adding 400 μL of cold extraction solvent (Acetonitrile: Methanol: Milli-Q Water; 40:40:20) to each sample, followed by three cycles of sonication (120 sec, power 60, frequency 37 kHz using an Elma Elmasonic P sonicator) and vortexing (240 sec). The samples were then centrifuged at 14 000 rpm for 10 min at +4 °C, after which the supernatants (350 µl) were transferred to evaporation tubes and gently evaporated dry under nitrogen stream. Dried samples were resuspended in 40 µl cold extraction solvent (Acetonitrile: Methanol: Milli-Q Water; 40:40:20) and transferred to HPLC auto sampler glass vials.

### LC-MS setup and data analysis

The analysis of the metabolites of interest was performed by injecting 2 μL sample extract to a Thermo Vanquish UHPLC coupled with a Q-Exactive Orbitrap quadrupole mass spectrometer equipped with a heated electrospray ionization (H-ESI) source probe (Thermo Fischer Scientific). Chromatographic separation of metabolites was done using a SeQuant ZIC-pHILIC (2.1 × 100 mm, 5 μm particle) column (Merck), which was maintained at +40 °C. Scanning was performed using full MS and polarity switching mode in mass range 55 to 825 m/z with the following settings: a resolution of 70,000, the spray voltages: 4250 V for positive and 3250 V for negative mode, the sheath gas: 25 arbitrary units (AU), and the auxiliary gas: 15 AU, sweep gas flow 0, capillary temperature: +275 °C, S-lens RF level: 50.0. Instrument control was operated with the Xcalibur 4.1.31.9 software (Thermo Fischer Scientific). The auto-sampler was used to perform partial loop with needle overfill injections for the samples at +5 °C. Gradient elution was carried out with a flow rate of 100 μL/min, and the total run time was 24 minutes per injection including a 2-minute equilibration step. The gradient elution started with 2 minutes 80 % mobile phase B (Acetonitrile), and then gradually increased to 80 % mobile phase A (20 mM ammonium hydrogen carbonate in water, adjusted to pH 9.4) until 17 minutes, after which the gradient switched back to 20 % mobile phase A at 17.1 minutes and equilibrated to the initial conditions for the last 7 minutes.

TraceFinder 5.1 software (Thermo Fischer Scientific) was used for data integration, the final peak integration, and peak area calculation of each metabolite. Peak smoothening was adjusted to 7, and the sample peak threshold was set to 50,000 m/z. The metabolite data was checked for peak quality (poor chromatograph), relative standard deviation (20% cutoff) and carry-over (20% cutoff). Data quality was monitored throughout the run by using a pooled sample as Quality Control (QC), which was prepared by combining an aliquot of 3 μL each study sample and injecting it after every 10th sample throughout the run.

### Metabolomics Data Analysis

Further, the data was explored and analyzed using Metaboanalyst site following the recommended settings. Metabolites with missing values in ≥20% of the samples were excluded from further analysis. Normalization by sum of sample peaks was used to normalize the data and log transformation (glog2) and autoscaling (mean-centred and divided by standard deviation of each variable) was used. Pathway enrichment analysis of metabolomics dataset was done for all increased and decreased metabolites of significance (p< 0.05) values using recommended settings of pathway analysis module in Metaboanalyst.

### RNA extraction

Total RNA from heart samples was extracted from snap frozen tissues in Trizol reagent (Invitrogen) and homogenized with Precellys Lysing kit -Tissue homogenizing CKMix (Bertin Technologies) and Precellys w-24 (Bertin Technologies). The RNA extracted was treated with DNAse and purified through RNeasy Qiagen minicolumns. The manufacturer’s instructions were followed for the extraction. The RNA quality was checked using Tapestation. Samples with a minimum of RIN value 7 were used for RNA-sequencing.

Library preparation and high-throughput sequencing were conducted at the Beijing Genomics Institute (BGI, Shenzhen, China) using BGI’s standard protocols. Sequencing was performed on the BGISEQ-500 platform with 100 bp paired-end (PE100) reads.Differential gene expression analysis was conducted in the R environment using the DESeq2 package^62^ with default parameters. A negative binomial generalized linear model and Wald test were applied to estimate p-values. Low-expression outliers were excluded based on Cook’s distance to enhance p-value adjustment accuracy. Multiple hypothesis testing corrections were performed using the Benjamini–Hochberg procedure.

### qPCR

RNA extracted from heart tissues were DNase-treated RNA (normalized across samples) was used for cDNA synthesis using the Maxima first-strand cDNA synthesis kit (Thermo Fisher Scientific) before qPCR using SensiFAST SYBR No-ROX kit (Bioline) and primers (details in Supplementary Table) according to the manufacturer’s instructions. The amplification level of the assayed gene (6 technical replicates per control and mutator mice) was normalized to *ACTB* and analyzed.

### Quantification and Statistical Analysis

Statistical analyses and their graphical representation were performed in GraphPad Prism v.7.0 software (GraphPad Software, USA). Statistical test used in each experiment is indicated in figure legend. P-values of less than 0.05 were considered statistically significant. Outlier analysis for all analyses was done using GraphPad Prism 7.0 ROUT method (Q=1%) and false positive analysis for metabolomics with Benjamin-Hochberg method, critical value of 0.2.

## Acknowledgments

The authors wish to thank Markus Innilä, Babette Hollman,Tuula Manninen, and Helena Puro for technical support in this work; Biomedicum Flowcytrometry Unit, University of Helsinki, Animal hospital at University of Helsinki for blood count analysis, Metabolomics core facility at university of Helsinki for its services, Helsinki University Laboratory Animal Centre for excellent maintenance of the animals. We thank Pekka Katajisto for providing antibodies used for FACS analysis from bone marrow cells.

Authors wish to acknowledge and thank Niagen Bioscience, the parent company of ChromaDex for providing NR for this study.

## Author Contributions

N.A.K developed the concept, tested hypotheses, designed and performed the experiments, analyzed data, interpreted results, supervised overall project and wrote and edited the manuscript. K.A performed experiments, analysed and interpreted FACS data of bone marrow and Peripheral blood. L.E performed redox metabolite measurement and performed experiments, commenting on the manuscript. S.P, J.L. performed experiments and analyzed data. C.J performed Oroboros experiments from mouse bone marrow cells. A.Z performed the bioinformatic analysis of RNA seq data. R.K performed an echo for the mouse heart function and interpreted the data. A.S. supervised the overall project, interpreted results,wrote and edited the manuscript.

