## Supplemental Fig 1-4 for "NAD^+^ precursor treatment prevents cardiomyopathy but disrupts erythroid maturation in mitochondrial progeria"

Supplementary figure 1

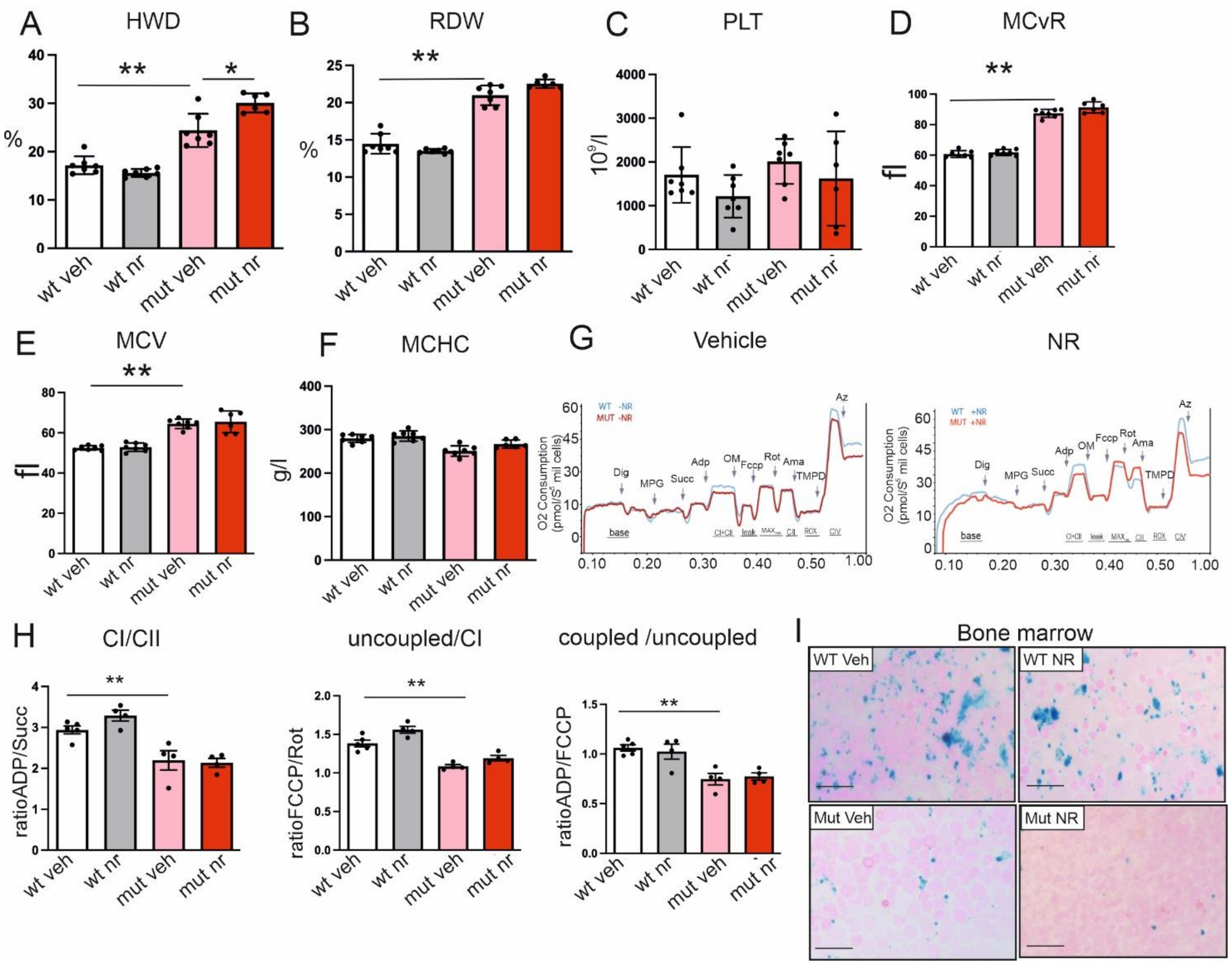


**Supplementary Figure 1 Mutators show decreased CI activity and lower iron content in bone marrow . related to Figure 1**

(A) Hemoglobin Distribution Width (HDW) displaying the variation in hemoglobin content within red blood cells in peripheral blood (n = 6 mice per group).

(B) Red Cell Distribution Width (RDW) in red blood cell size in peripheral blood (n = 6 mice per group).

(C) Platelet Counts in peripheral blood mice (n = 6 mice per group)

(D) Mean reticulocyte volume (McVR), quantification of average reticulocyte size in peripheral blood. (n = 6 mice per group)

(E) Mean Corpuscular Volume (MCV), average volume of red blood cells in peripheral blood (n = 6 mice per group).

(F) Mean Corpuscular Hemoglobin Concentration (MCHC), concentration of hemoglobin to the volume of red blood cells in peripheral blood (n = 6 mice per group).

(G) Respiration traces in bone marrow cells of mutator and control mice with vehicle and NR treatment, C = coupled, UC = uncoupled (n = 4 mice per group).

(H) Complex I activity and respiration in bone marrow, uncoupled versus coupled respiration in bone marrow, Complex I-dependent respiration in bone marrow (n = 4 mice per group)

(I) Iron (Fe) Staining (blue) of bone marrow cells to evaluate iron deposition and distribution. Scale bars represent 100 µm.

Statistical analyses were performed using Student’s t-test or two-way ANOVA as appropriate. *p ≤ 0.05, **p ≤ 0.01, ***p ≤ 0.001, ****p ≤ 0.0001. Data are expressed as mean ± SEM.

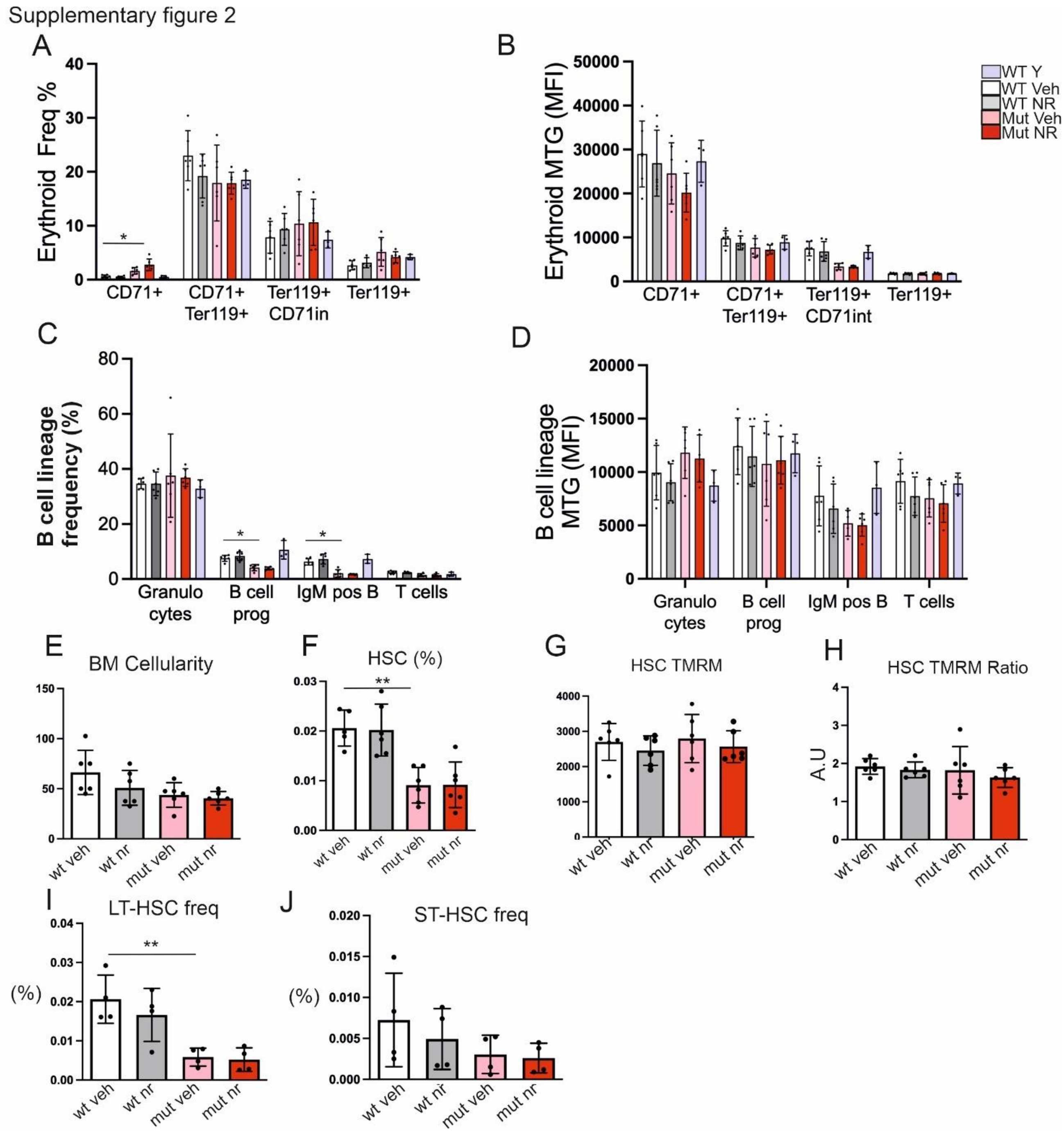


**Supplementary Figure 2: NR treatment did not change cell types in both mutator and wt mice related to figure 1 and figure 2**

(A) Erythroid Progenitors Frequency: Flow cytometry analysis showing the frequency of erythroid progenitors in bone marrow or mutator and control mice (n = 6 mice per group, young wt , n=3).

(B) Erythroid MitoTracker Green (MTG) Staining: Flow cytometry analysis showing the percentage of MitoTracker Green-positive erythroid cells in peripheral (n = 6 mice per group, young wt , n=3).

(C) B cell lineage frequency: Flow cytometry analysis showing the frequency of B cell lineage cells in bone marrow or mutator and control mice (n = 6 mice per group, young wt , n=3).

(D) MitoTracker Green (MTG) Staining: Flow cytometry analysis showing the percentage of MitoTracker Green-positive B cell lineage cell in mutator and control mice (n = 6/ grp, young wt , n=3).

(E) Bone Marrow Cellularity: Bar graph depicting the total cellularity of bone marrow in progeria mice (n = 6 mice per group).

(F) HSC % in total bone marrow cells.

(G-H) TMRM intensity in hematopoietic stem cells (HSCs) from bone marrow (n = 6 mice per group), reflecting mitochondrial membrane potential and HSC TMRM ratio in bone marrow cells.
(I) Long-Term Hematopoietic Stem Cell (LT-HSC) Frequency in bone marrow (n = 4 mice per group).
(J)Short-Term Hematopoietic Stem Cell (ST-HSC) Frequency in bone marrow (n = 4 mice per group).
(Statistical analyses were performed using Student’s t-test or two-way ANOVA as appropriate. *p ≤ 0.05, **p ≤ 0.01, ***p ≤ 0.001, ****p ≤ 0.0001. Data are expressed as mean ± SEM.

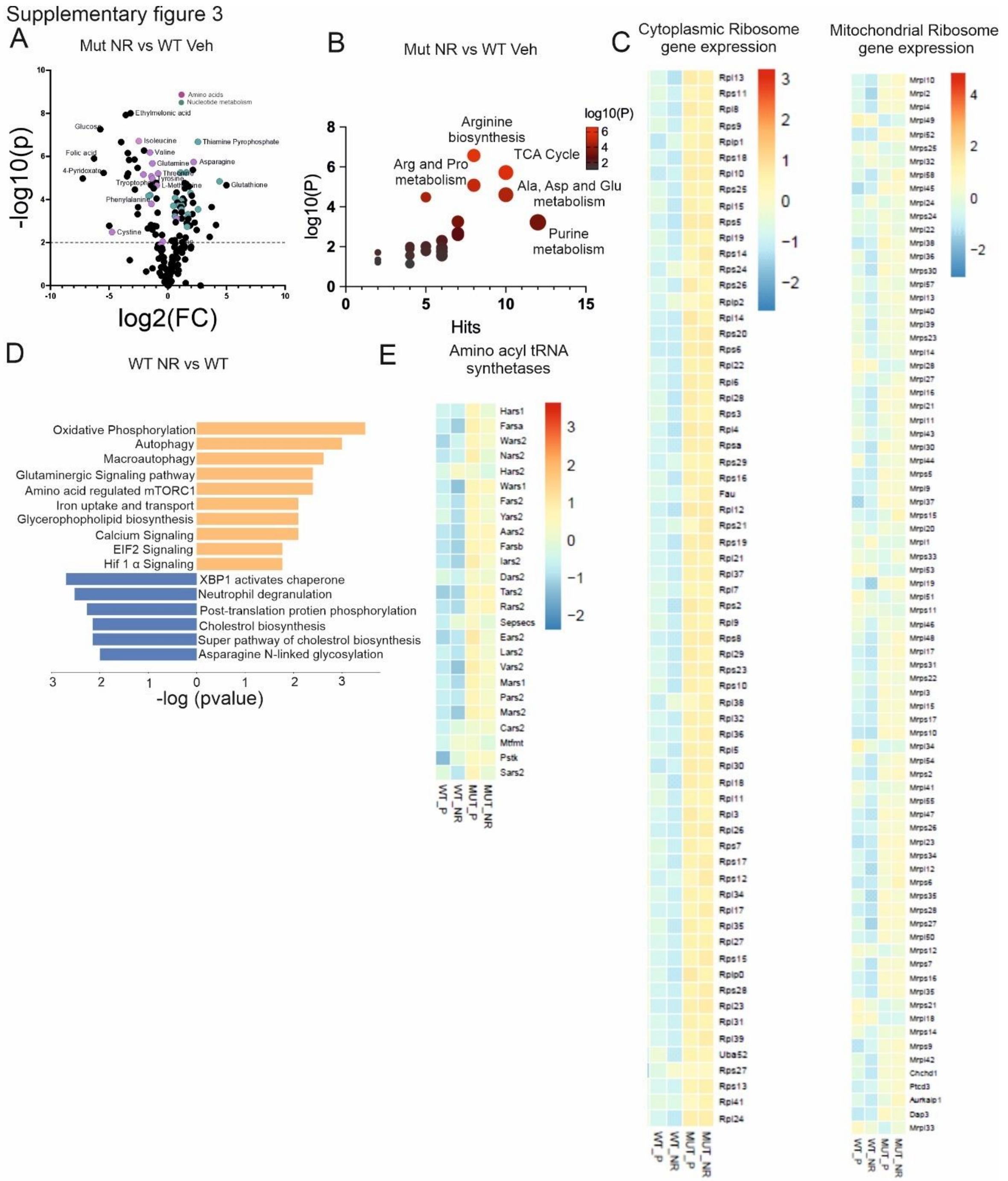


**Supplementary Figure 3 Mutator bone marrow cells show dysregulated mitochondrial and cytosolic protein synthesis pathway related to Figure 2 and Figure 3**
(A) Volcano plot depicting significantly changed metabolites in NR-treated mutator bone marrow compared to vehicle control (n = 5 per group).
(B) Targeted pathway analysis for the top metabolites in the bone marrow of NR treated mutator mice compared to vehicle control (n = 5 per group).

(C) Heatmap showing mitochondrial and cytoplasmic ribosomal gene expression in mutator mice bone marrow compared to control as group average (WT Veh: n = 4, WT NR: n = 5, Mut Veh: n = 7, Mut NR: n = 7).
(D) Transcriptomic pathways changed in the bone marrow transcriptome in NR-treated control mice compared to vehicle-treated control mice by ingenuity pathway analysis (WT Veh: n = 4, WT NR: n = 5)
(E) Heatmap showing amino acyl tRNA synthetase gene expression in mutator mice bone marrow compared to control as group average (WT Veh: n = 4, WT NR: n = 5, Mut Veh: n = 7, Mut NR: n = 7).

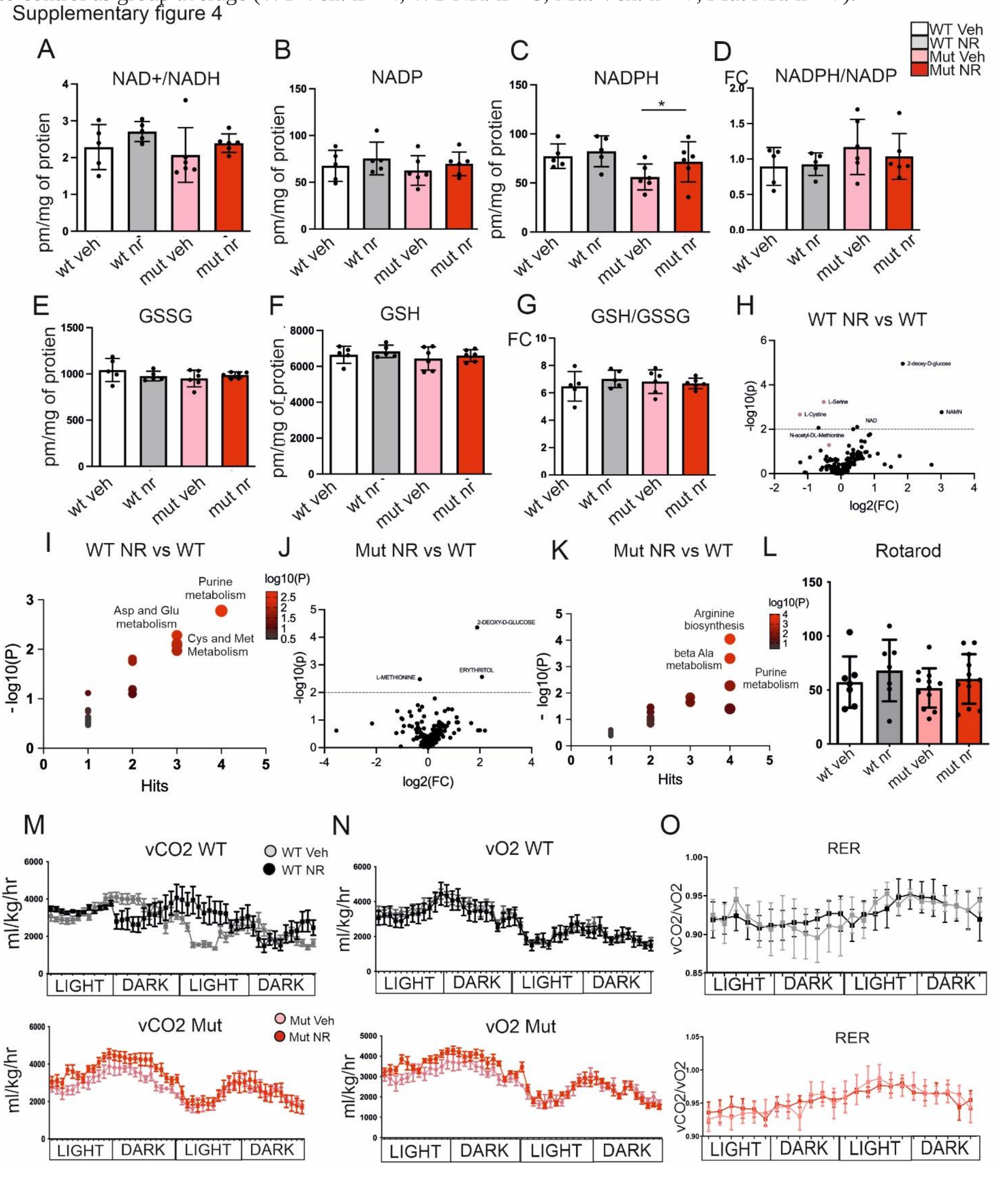


**Supplementary Figure 4: NR treatment preserves redox balance in heart and mildly improves whole body metabolism. related to figure 4.**
(A) Ratio of NAD+/NADH in heart tissue of mutator and WT mice with vehicle or NR treatment (n = 6 mice per group).

(B-D) NADP, NADPH levels and NADPH/$NADP^+$ Ratio in heart tissue of mutator and WT mice with vehicle or NR treatment (n = 6 mice per group).
(E-G) GSSG, GSH levels and GSH/GSSG Ratio in heart tissue of mutator and WT mice with vehicle or NR treatment (n = 6 mice per group).
(H) Volcano plot showing significantly changed metabolites in NR-treated control heart compared to vehicle-treated control (n = 5).
(I) Significantly changed metabolite pathway for the top metabolites top metabolites in NR-treated control heart compared to vehicle-treated control (n = 5).
(J) Volcano plot showing significantly changed metabolites in NR-treated mutator heart compared to vehicle-treated control (n = 5).
(K) Significantly changed metabolite pathway for the top metabolites in NR-treated mutator heart compared to vehicle-treated control (n = 5).
(L)Rotarod Performance: Bar graph showing the latency to fall from the rotarod in wild-type mice (WT Veh: n = 7, WT NR: n = 7, Mut Veh: n = 12, Mut NR: n = 12).
(M) Whole body metabolism analysis; metabolic cages. Carbon dioxide production recorded for 24h in room temperature mutator and control mice with vehicle and NR diet (n = 6 mice per group). Light=inactivity. Dark=active period.
(N) Whole body metabolism analysis; metabolic cages. Oxygen consumption recorded for 24h in room temperature mutator and control mice with vehicle and NR diet (n = 6 mice per group). Light=inactivity. Dark=active period.
(O) Respiratory exchange ratio (RER, $vCO_2/vO_2$) in mutator and control mice with vehicle and NR diet (n = 6 mice per group). RER ~ 1, preferred fuel carbohydrates; RER ~ 0.7, lipids as fuel. Light=inactivity. Dark=active period.

Statistical analyses were performed using Student's t-test or two-way ANOVA as appropriate. *$p \leq 0.05$, **$p \leq 0.01$, ***$p \leq 0.001$, ****$p \leq 0.0001$. Data are expressed as mean ± SEM.
